# Yield potential definition of the chilling requirement reveals likely underestimation of the risk of climate change on winter chill accumulation

**DOI:** 10.1101/285361

**Authors:** José Antonio Campoy, Rebecca Darbyshire, Elisabeth Dirlewanger, José Quero-García, Bénédicte Wenden

## Abstract

Evaluation of chilling requirements of cultivars of temperate fruit trees provides key information to assess regional suitability, according to winter chill, for both industry expansion and ongoing profitability as climate change progresses. Traditional methods for calculating chilling requirements use climate controlled chambers and define chilling requirements using a fixed bud burst percentage, usually close to 50% (CR-50%). However, this CR-50% definition may estimate chilling requirements that lead to flowering percentages that are lower than required for orchards to be commercially viable. We used sweet cherry to analyse the traditional method for calculating chilling requirements (CR-50%) and compared the results with a more restrictive method, where the chilling requirement was defined by a 90% bud break level (CR_m_-90%). For sweet cherry, this higher requirement of flowering success (90% as opposed to 50%) better represents grower production needs as a greater number of flowers lead to greater potential yield. To investigate the future risk of insufficient chill based on alternate calculations of the chilling requirement, climate projections of winter chill suitability across Europe were calculated using CR-50% and CR_m_-90%. Regional suitability across the landscape was highly dependent on the method used to define chilling requirements and differences were found for both cold and mild winter areas. Our results suggest that bud break percentage levels used in the assessment of chilling requirements for sweet cherry influence production risks of current and future production areas. The use of traditional methods to determine chilling requirements can result in an underestimation of productivity chilling requirements for tree crops like sweet cherry which rely on a high conversion of flowers to mature fruit to obtain profitable yields. This underestimation may have negative consequences for the fruit industry as climate change advances with climate risk underestimated.

## Introduction

Temperate deciduous fruit trees represent an important economic and food resource. Maintaining and expanding temperate fruit production is an industry priority and is likely to become more challenging under anthropogenic climate change. One aspect of tree physiology which will likely be influenced by climate change is sufficient accumulation of winter chilling (Luedeling et al. 2011). In temperate fruit trees, the timing of bud break and flowering is highly correlated with a proper release of endodormancy, or rest, which is controlled by the exposure to cold temperatures in winter (Saure 1985). During this rest period a certain exposure to cold temperatures, defined as the chilling requirement, must be satisfied prior to the bud being released from endodormancy (Lang et al. 1987). This period of rest prevents bud break in response to conditions suitable for growth conditions, such as a warm spell in winter, which would then expose sensitive tissues to subsequent damaging cold conditions. The consequences of insufficient chill accumulation have been extensively studied and include delayed, light, irregular and prolonged bud break and flowering (Samish 1953; Erez and Couvillon 1987; Erez 2000) as well as damage to bud development including deformed buds (Petri and Leite 2004; Viti et al. 2008). Within the context of climate change, these sub-optimal production outcomes may increase in frequency (Guy 2014). To predict such potential impacts to assist with orchard climate adaptation and guide breeding strategies, evaluation of cultivar specific chilling requirements (CR) is needed (Dennis 2003; Campoy et al. 2011; Chuine et al. 2016).

Precise determination of the CR is not possible under field conditions. As such, various experimental methods have been used to evaluate dormancy depth (Cook et al. 2017; Vitra et al. 2017), date of dormancy release (Dennis 2003; Dantec et al. 2014; Castède et al. 2014; Chmielewski and Götz 2016) and to assess the relationship between bud break and chilling and forcing temperatures (Harrington et al. 2010; Luedeling and Gassner 2012; Andreini et al. 2014; Laube et al. 2014). Forcing experiments using controlled climate chambers have been commonly used to assess bud dormancy progression since the method was first proposed for the evaluation of CR in peach (Bennet 1949; Weinberger 1950). This approach involves sampling tree cuttings through autumn and winter and evaluating bud break after they have been exposed to warm temperatures in a controlled climate chamber (Tabuenca 1967; Dennis 2003; Vitasse and Basler 2014). In implementing this methodology two main criteria have been used to determine CR: 1) percentage of floral or vegetative bud break within a given period of time in the chamber (e.g. 50% break after 10 days) and 2) the time required in the forcing chamber to reach a given stage of development (e.g. bud break) (Saure 1985; Dennis 2003). the first approach often determines CR as the chill accumulation in the field leading to 30-50% bud break after 7 to 10 days in the forcing chamber (Hauagge and Cummins 1991; Ruiz et al. 2007; Viti et al. 2010; Campoy et al. 2012; Castède et al. 2014). This criteria varies by study with for instance, CR being determined after the emergence of only 3-4 flowers in the forcing chamber (Chmielewski and Götz 2016; Götz et al. 2017).

These criteria may not adequately represent CR that are required for profitable fruit production, in particular for crops which require high conversion of floral buds to fruit (e.g. sweet cherry). These businesses rely on high and consistent bud break rates which set a high yield potential. Poor rates of floral and vegetative bud break, that is the proportion of floral and vegetative buds that truly open, can have a major impact on temperate fruit crop productivity (Erez 2000). For small fruits, such as sweet cherry, a large number of viable flowers is required to reach commercial productivity, with a maximum bloom level ideal. Therefore, setting a CR value which only leads to 30-50% bud break may underestimate the CR needed for commercial profitability. This in turn will misjudge the potential risk of climate change on cherry production.

In this work, one of the most commonly employed methods in temperate fruit trees to estimate CR (chilling accumulation in field that leads to 50% bud break after 10 days forcing) was used for sweet cherry and was contrasted with a modified method that sets a higher bud break percentage to better match requirements for high yield potential (90% bud break). We used CR determined from both methods to create climate change projections of the potential risk of insufficient chill accumulation for sweet cherry across Europe. This work emphasises the need to evaluate CR in terms of the minimum requirement of chill for commercial viability to best prepare industry and breeding programs for expansion and climate change.

## Materials and methods

### Bud break observations

We chose four sweet cherry cultivars for analysis encapsulating a presumed range in CR based on flowering dates (Fig. 1). The trees used for the experiments were grown following commercial orchard management in the fruit tree experimental orchards of the Institut National de la Recherche Agronomique (INRA)-Bordeaux research centre, located in Bourran, in Lot-et-Garonne (France, 44°19’N 0°24’E, 70m asl) and 60 km away in Toulenne, in Gironde (France, 44°34’N 0°16’W, 8m asl). Cultivars ‘Cristobalina’ and ‘Regina’ were grown in Bourran, ‘Fertard’ in Toulenne and ‘Burlat’ in both sites. Dates for beginning of flowering (5-10% open flowers, BBCH 61; Fadón et al., 2015) and maturity (BBCH 89) were recorded for the four cultivars between 2000 and 2017 in order to define the differences among cultivars. For the experimental data, two or three branches bearing floral buds were randomly cut from the two trees per cultivar and placed in a climate-controlled chamber under forcing conditions (25°C, 16h light/8h dark) each fortnight between 1^st^ September and 8^th^ April (2015-2016 and 2016-2017 seasons), except for ‘Cristobalina’ which was sampled only in the 2015-2016 season.,. Every two or three days the total number of floral buds that reached bud break (BBCH stage 53; Fadón et al. 2015) was recorded. Bud break percentage was calculated as the percentage of flower buds at BBCH stage 53 in relation to the total number of floral buds on the branches.

**Fig. 1.**
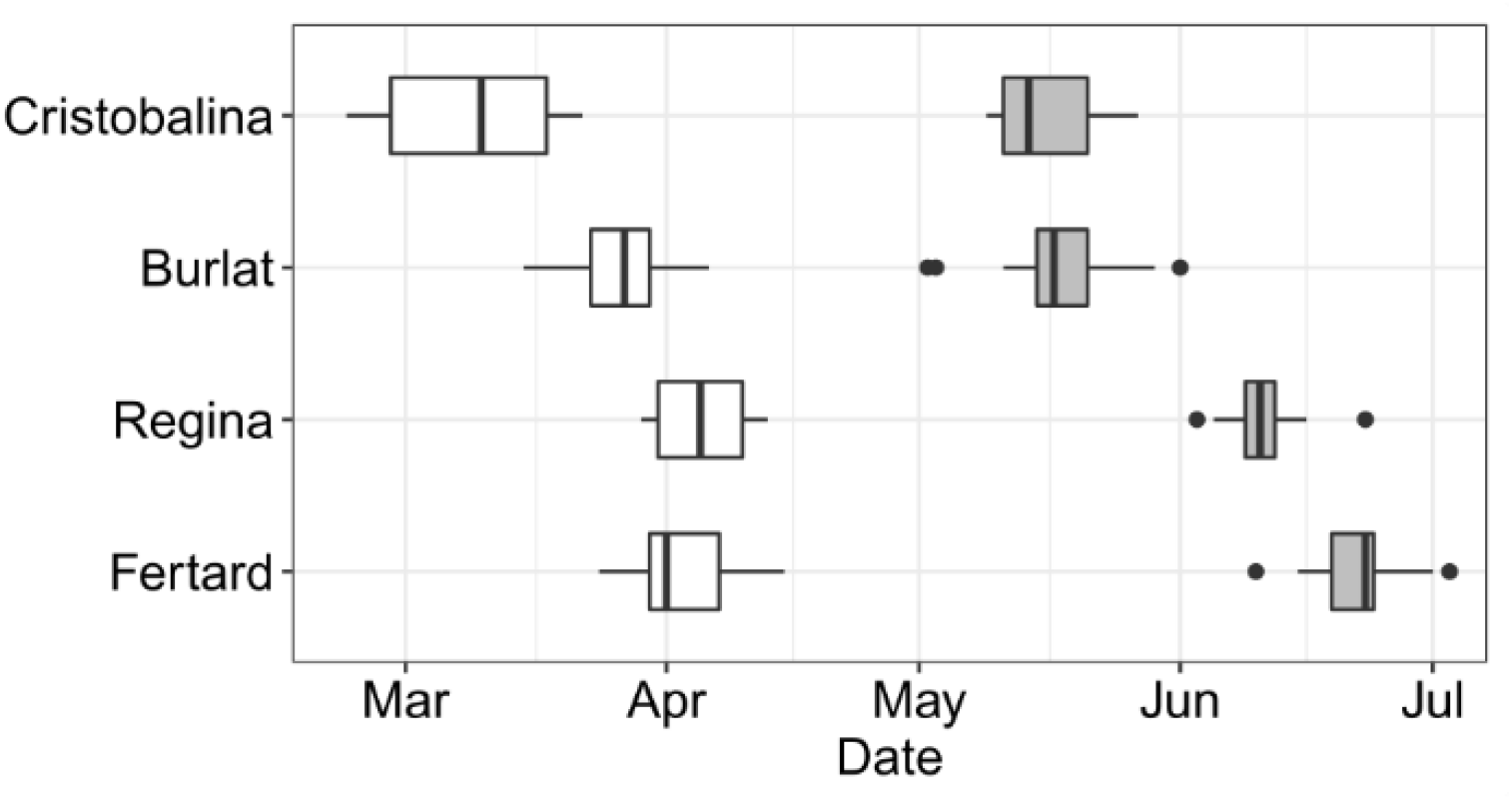
Range of flowering (white boxes) and maturity (grey boxes) dates for the four studied cultivars (2000 – 2017)

### Chill requirement estimation

Chill was calculated in chill portion (CP) using the Dynamic chill model (Fishman et al. 1987), which has previously been found to be the best fitting chill model among several tested models for a range of fruit tree species (Alburquerque et al. 2008; Luedeling et al. 2009b, 2013a; Campoy et al. 2011; Guo et al. 2013).

To calculate CP hourly, we collected mean temperatures from local weather stations (years 2015-2017, INRA CLIMATIK, https://intranet.inra.fr/climatik, stations 47038002 and 33533002 for Bourran and Toulenne, respectively). CP were calculated using the original model parameters provided in the R package chillR (Luedeling et al. 2013b; Luedeling 2018). Chill was accumulated from October 1^st^ until the sampling date.

To assess CR, we used two methods. Firstly, we applied the commonly used approach which defines CR as the accumulated chill in the field which leads to 50% of the buds surpassing stage BBCH 55 after 10 days in forcing conditions (CR-50%) (Ruiz et al. 2007; Alburquerque et al. 2008; Campoy et al. 2012; Sánchez-Pérez et al. 2012; Azizi Gannouni et al. 2017). Secondly, we used a similar approach but with a longer period in forcing conditions (2-3 weeks) and a higher bud break percentage, 90%, required to define CR (CR_m_-90%).

### Future climate data

For the analyses of future climate risk, we used two gridded datasets of temperatures in Europe. These were regionally downscaled climate datasets for historical and projected temperature made available by EURO-CORDEX (http://www.euro-cordex.net; Jacob et al., 2014). This dataset used the SMHI-RCA4 regional change model (50 km resolution, EUR-44) with projection data from the climate models MOHC-HadGEM2-ES and NOAA-GFDL-GFDL-ESM2M. Two scenarios (RCP 4.5 and RCP 8.5) were assessed for each climate model. We corrected the differences for the temperatures between EURO-CORDEX projections and recorded data using the European observation (E-OBS) gridded temperature dataset with 0.25° resolution (v.14; Haylock et al., 2008). For the bias correction, we calculated the average anomaly grid as the mean temperature difference between the E-OBS and regional climate model datasets for 1978-2005. Gridded annual chill accumulation values were then calculated for the datasets (E-OBS, MOHC-HadGEM2_ES and NOAA-GFDL-GFDL-ESM2M), after adding the anomaly grid to the RCM datasets.

Hourly temperatures were generated for the three climate datasets using the R package chillR (Luedeling 2018) which uses a sine curve and a logarithmic function for day-time and night-time temperatures, respectively. Chill was then accumulated between October 1^st^ and March 31^st^.

Safe winter chill (SWC; Luedeling et al., 2009b) was used to estimate future production risk in Europe based on winter chilling conditions for each cultivar and for the two CR estimated using the two methods. SWC is the amount of winter chill that can be reliably expected in 90% of all years, or the 10^th^ percentile of the dataset. Here this corresponds to the data within each of the time periods analysed (1975-2005, 2026-2050, 2075-2100). This metric is meaningful to fruit producers, because failure to meet chilling requirements in more than 10% of years is likely to render production uneconomical (Luedeling et al. 2009a).

## Results

### Bud break percentage based on chill accumulation

Evaluation of bud break response to accumulated chill in the field prior to cuttings revealed that low - but not nil - bud break percentages were observed for branches that had been sampled in the early season (mid-November; Fig. 2). These results suggest that early (‘Cristobalina’) and mid-flowering (‘Burlat’) cultivars could slightly respond (i.e. less than 5% bud burst) to forcing temperatures after accumulating only 13 CP. The rate of bud break percentage increased as more chill was accumulated in the field. Using ‘Burlat’ as an example, bud break percentage increase markedly faster for branches that accumulated at least 90 CP than for branches sampled earlier during chill accumulation (e.g. 32 CP). The maximum bud break percentage, illustrated by the plateau for each line, (Fig. 2) also increased with more chill accumulation. These results indicate that the amount of accumulated chill primarily drives bud break percentages as the buds were all exposed the same amount of time to forcing temperatures (Fig. 2).

**Fig. 2.**
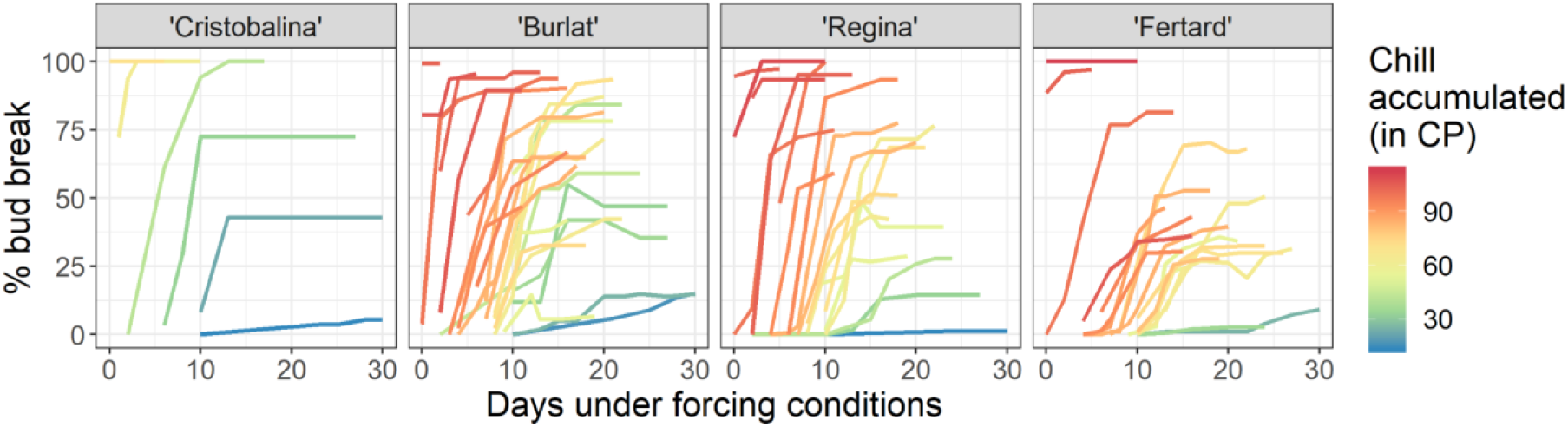
Floral bud break percentage for the four cultivars with various field chill accumulation values (in chill portions). Note that no results for ‘Cristobalina’ were recorded for chill accumulation greater than 69 CP as bud burst had begun in the field.

The response of bud break percentage to accumulated chill notably differ between cultivars.

For example, the early flowering cultivar ‘Cristobalina’ (i.e. low-chill cultivar) was found to have higher bud break potential (45% bud break) after low levels of accumulated chill (32 CP) whereas the late flowering (i.e. high-chill cultivar) cultivar ‘Regina’ only reached 14.5% bud break after accumulating 32 CP. For the late flowering cultivars to meet the equivalent 45% bud break, 53 CP and 65 CP were needed for ‘Regina’ and ‘Fertard’, respectively. We also observed this pattern for the capacity to reach high bud break percentage. After accumulating 40 CP, ‘Cristobalina’ and ‘Burlat’ reached 100% and 84% bud break, respectively, whereas the late flowering cultivars (‘Regina’ and ‘Fertard’) only reached 28% and 16% bud break, respectively (Fig. 3, Fig. S1).

**Fig. 3.**
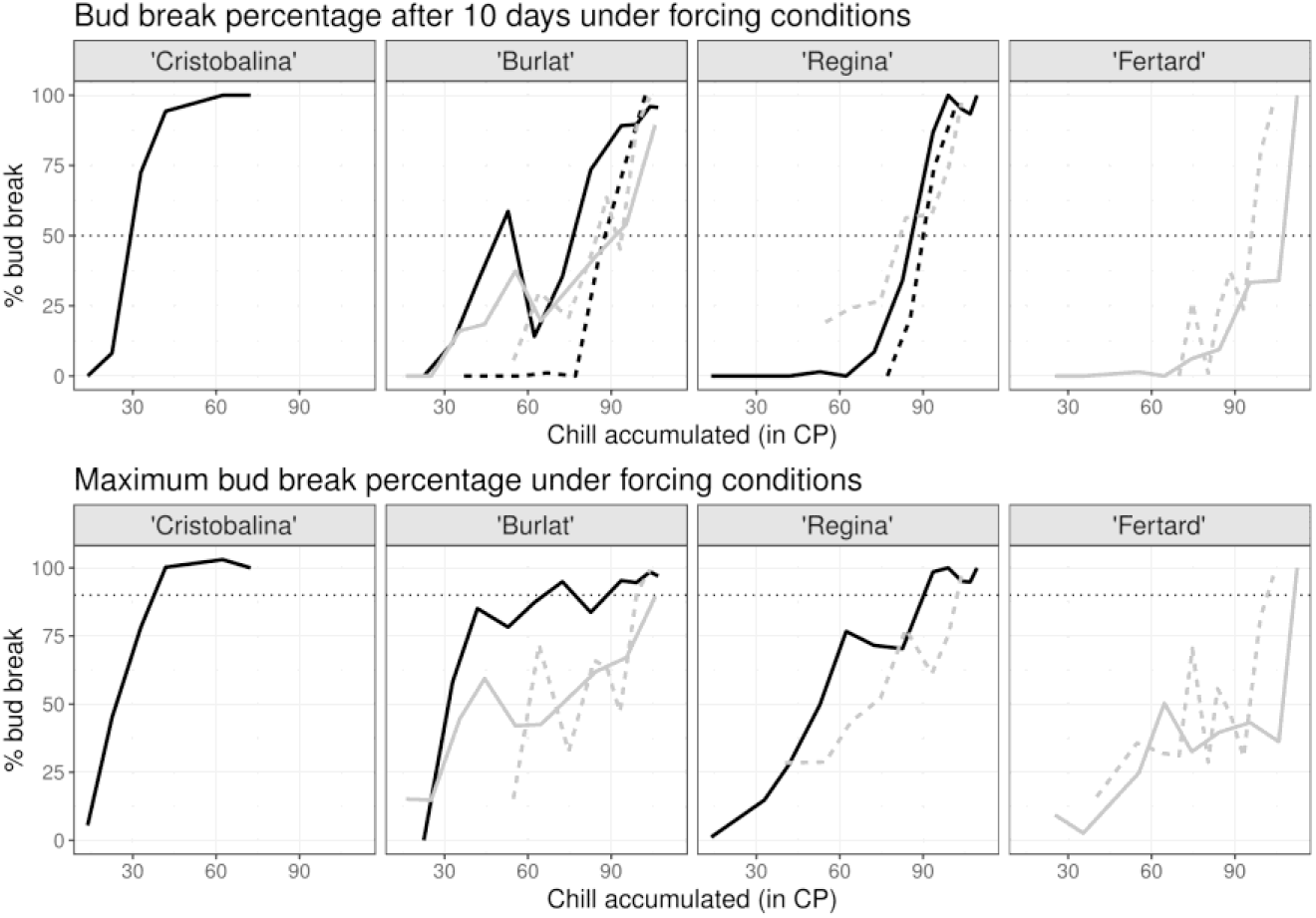
Bud break data used to calculate CR-50% and CR_m_-90%. Upper panel: bud break percentage after 10 days under forcing conditions; lower panel: maximal bud break percentage reached under forcing conditions. Horizontal dotted lines represent the 50% and 90% thresholds for CR-50% and CR_m_-90% respectively. Black and grey lines represent data for cultivars grown in Bourran and Toulenne, respectively. Solid lines are for 2015-2016 data while dashed lines are for 2016-2017 data

### Chilling requirement estimation

Chilling requirement values calculated using the common approach (CR-50%) were highly variable between years (Fig. 3, Table 1). The CR range calculated using a 90% bud break threshold (CR_m_-90%) were less variable.

**Table 1.**
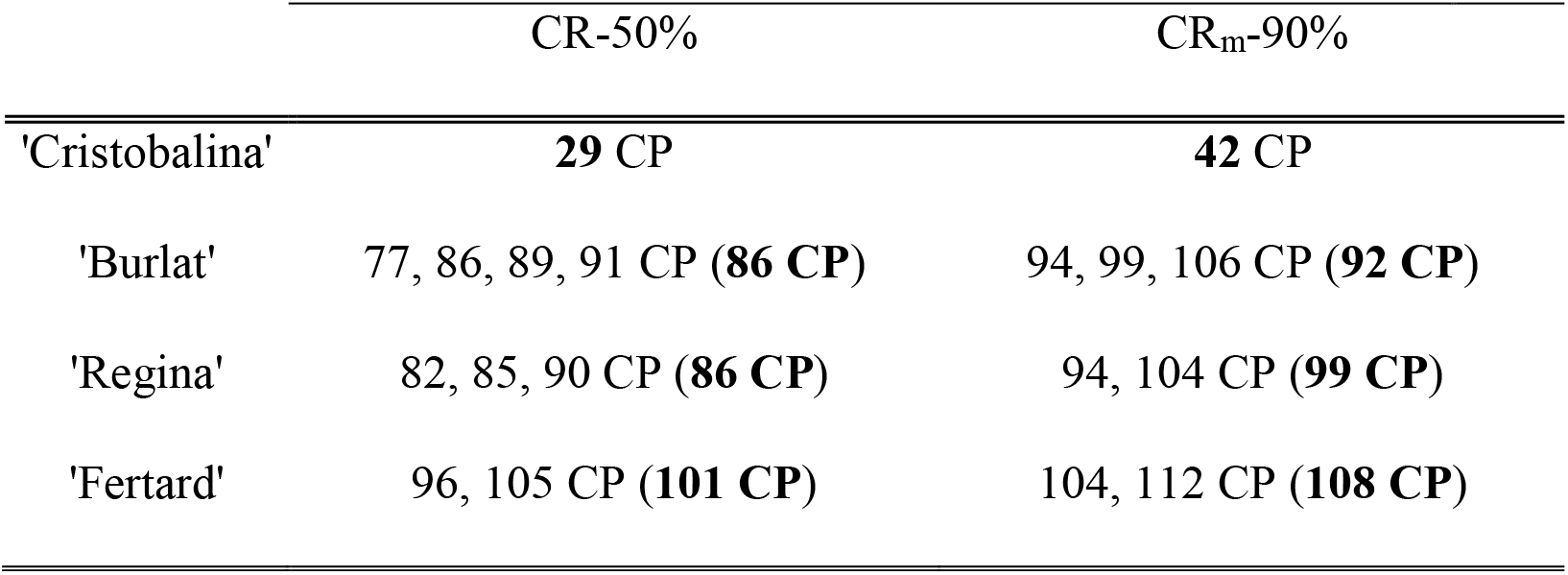
Chill requirement values (CP) between the four cultivars and the method used to calculate CR. Bold values are the mean CR recorded.

|  | CR-50% | CR<sub>m</sub>-90% |
| --- | --- | --- |
| 'Cristobalina' | **29 CP** | **42 CP** |
| 'Burlat' | 77, 86, 89, 91 CP (**86 CP**) | 94, 99, 106 CP (**92 CP**) |
| 'Regina' | 82, 85, 90 CP (**86 CP**) | 94, 104 CP (**99 CP**) |
| 'Fertard' | 96, 105 CP (**101 CP**) | 104, 112 CP (**108 CP**) |

Taking the average chill accumulation to reach 50% and 90% bud break, CR for each cultivar were estimated (Table 1). CR differs with cultivar and by the method used to evaluate it but the CR estimated by CR_m_-90% are all higher than CR-50%.

### Future risk due to insufficient chill

Estimation of the risk of insufficient chill accumulation across Europe based on the two methods to determine CR (Table 1) exposed areas potentially unsuitable for the cultivation of each sweet cherry cultivar (Figs 4, 5, S2 and S3). We focused on MOHC-HadGEM2-ES, rcp8.5, projecting the highest increase in temperatures (most pessimistic; Fig. 4) and NOAA-GFDL-GFDL-ESM2M, rcp4.5, projecting the lowest increase in temperatures (optimistic; Fig. 5). Areas defined in Figures 4 and 5 reveal that for the four cultivars, the regions at risk of not meeting CR increase substantially by 2075-2100 using both estimations of CR (CR-50% and CR_m_-90%). The area at risk is larger for CR_m_-90% than for CR-50% for both scenarios. Areas coloured in pink, show the areas that meet CR according to CR-50% but not according to CR_m_-90% (Figs 4 and 5). Therefore, these pink areas show the difference of estimated risk of insufficient chill depending on the method used to define CR. The difference in risk based on CR methodology is particularly evident for the high-chill cultivar ‘Regina’. Under both climate scenarios, a large area of Europe will not meet CR_m_-90% whereas using CR-50% a much smaller area of risk was found (Figs 4 and 5).

**Fig. 4.**
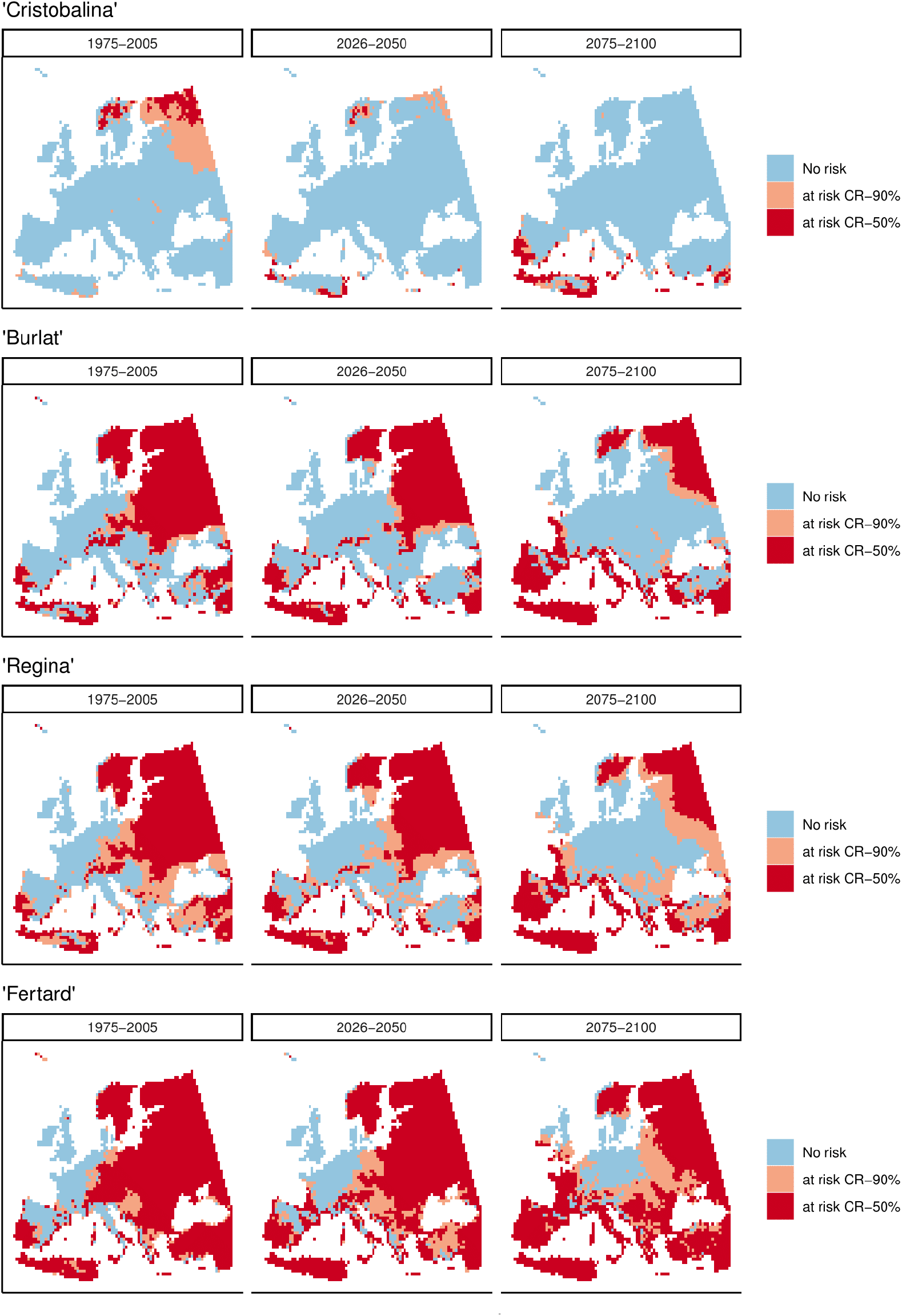
Maps of Europe showing where safe winter chill (10^th^ percentile) will not meet chilling requirements as estimated by forcing experiments (CR-50%; red areas) and by optimal bud break rate (CR_m_-90%; pink and red areas) for the four cultivars. Blue indicates CR will be met for both CR values. Thresholds for the chilling requirements calculated in Table1 were used. Scenario MOHC-HadGEM2-ES, rcp8.5, projects the highest increase in temperatures (most pessimistic)

**Fig. 5.**
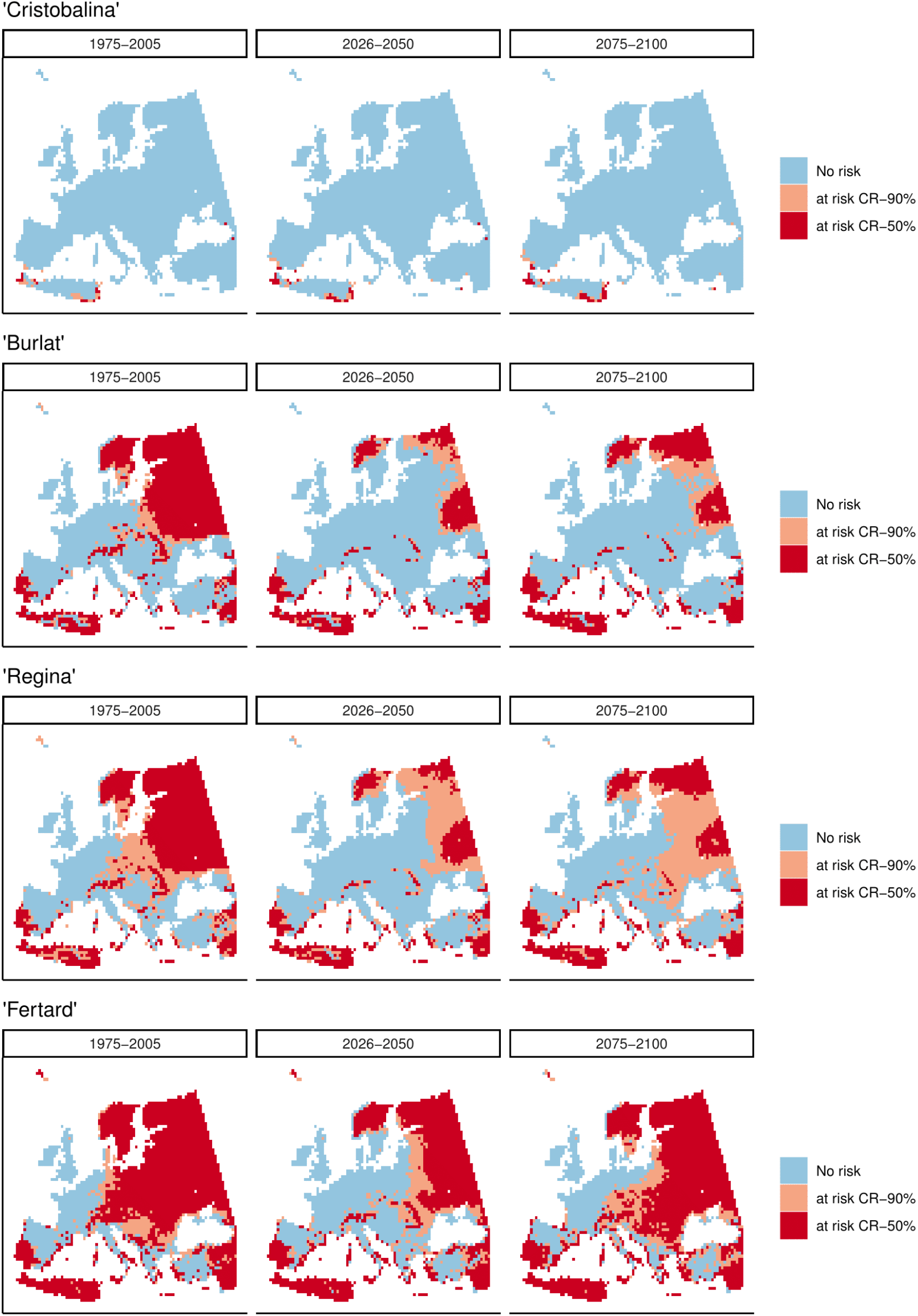
Maps of Europe showing where safe winter chill (10th percentile) will not meet chilling requirements as estimated by forcing experiments (CR-50%; red areas) and by optimal bud break rate (CRm-90%; pink and red areas) for the four cultivars. Blue indicates CR will be met for both CR values. Thresholds for the chilling requirements calculated in Table1 were used. Scenario NOAA-GFDL-GFDL-ESM2M, rcp4.5, projects the lowest increase in temperatures (most optimistic)

## Discussion

### Bud break percentage in response to chill accumulation

We evaluated bud break responses to various levels of chill accumulation in the field using forcing experiments on cultivars with contrasting chill requirements. The rate of bud break gradually increased with accumulated chill in the field, thus confirming that chill primarily drives the capacity of the buds to burst. Greater chill accumulation increased the maximum bud break percentage but with a different maximum response between cultivars (Fig. 3). Overall, the results indicate that the amount of accumulated chill can substantially limit the percentage of bud break even under optimal growth conditions (Fig. S1). This is especially true for the early flowering cultivar ‘Cristobalina’ which shows a very strict control of bud break percentage by the amount of chill accumulated with limited effect of heat accumulation (Fig. 4; Fig. S1). In addition, according to our data, there is a limit to the effect of heat on the bud break percentage. When bud break percentages were considered as a response to chill and heat accumulation (Fig. S1), results suggest that for a given chill accumulation bud break might never reach 100%. For example, bud break percentage in ‘Regina’ fails to reach 60% if chill accumulation is under 60 CP regardless of the level of heat accumulated. This provides a clear chill requirement limit on maximum bud break percentage.

The interaction of chilling with heat requirements and bud break was also shown in different temperate *Rosaceae*: cherry (Measham et al. 2017), peach (Couvillon and Erez 1985; Okie and Blackburn 2011), sour cherry (Felker and Robitaille 1985), apple (Powell 1986) and pear (Spiegel-Roy and Halston 1979); and in a notable group of forest species (Laube et al. 2014). This interaction has been associated to a ‘residual effect of dormancy’ (Erez 2000), a parameter indicating the distance to optimal chilling, and therefore, productivity (Okie and Blackburn 2011). This term is used when crops show symptoms of insufficient chilling (e.g. light bud break or fruit set and uneven foliation) even after satisfaction of their CR, usually corresponding to CR-30% or CR-50% (Campoy et al. 2011). Therefore, increasing the required bud break percentage when assessing CR will provide a more reliable measure of CR which should minimise productivity problems associated with the ‘residual effect of dormancy’ for new growing areas and improve commercial viability. Other studies that have calculated CR for species that set yield potential based on flower numbers (e.g. almond; Egea et al. 2003), may also consider recalculation of CR based on a commercial bud break requirement to assist with current and future industry planning.

### Chilling requirement calculation

The chill accumulation necessary to reach 90% (CR_m_-90%) of flower bud break was found to be higher than the chilling requirements estimated with the common forcing method (CR-50%) as shown in Table 1. This reflects that the traditional assessment of chilling requirements may underestimate the CR needed to reach commercial productivity in sweet cherry. It would not be advisable to choose CR-100% (100% bud break in the controlled environments) as a small percentage (i.e. 10%) allows for a buffer for minor physiological problems or accidental bud drop during the manipulation of plant material on growth chambers.

The potential underestimation of CR due to experimental design may be trivial for temperate areas characterised by cold winters, where chill accumulation in the field normally exceeds the CR of all cultivars with both CR-50% and CR_m_-90% easily achieved. For locations with more marginal chill accumulation, assessments of suitability based on CR-50% instead of CR_m_-90% will likely have a more significant impact. The underestimation of CR could lead to unexpectedly higher frequency and severity of productivity problems due to insufficient chilling fulfilment. This will be particularly applicable to low-chill cultivars that are grown in mild winter climates which are usually oriented to the profitable early-market (Egea et al. 2010).

Values for both CR-50% and CR_m_-90% were highly variable between years and sites, although less variable for CR_m_-90%. Similar variability was been reported for sweet cherry in Australia (Measham et al. 2017) and apricot in Spain and South Africa (Campoy et al. 2012). This variability in CR estimation should be appreciated when considering production risk. One potential source of variability in recorded CR is the chill model that we used in the analysis. Indeed, the Dynamic model, used here to quantify chilling, was developed for Mediterranean regions, and therefore may not be suitable for colder climates. However, past studies have shown that, regardless of the climatic conditions, the Dynamic model is currently the most accurate model to use for predicting chill requirements (Luedeling et al. 2011; Luedeling 2012). Further investigation will be necessary to evaluate the accuracy of the model to ensure the robustness of our analysis and predictions. This line of enquiry would similarly benefit other assessments of CR in small fruit temperate species such as sweet cherry (Alburquerque et al. 2008) or almond (Egea et al. 2003) and projections of winter chill (Luedeling et al. 2011). It would also be valuable to better understand the source of plasticity in bud responses to cold in order to improve chill models for contrasted climatic conditions.

### Future production risk due to insufficient chill

The projections of chill accumulation provide new evidence of climate change risks for the cultivation of sweet cherry across Europe. Risk of insufficient chill accumulation increased with the chilling requirement of the cultivar and the method used to calculate it (CR-50% or CR_m_-90%). In the Mediterranean basin (a marginal chill area), only extra-low chill cultivars, such as ‘Cristobalina’, were found suitable across the 21^st^ century. However, for high-chill cultivars, suitable areas were restricted and almost the whole of the continental Europe recorded risk, especially using the CR_m_-90% estimation (Figs. 4 and 5). These results highlighting CR underestimation compounded by warming winters provide a warning message concerning the sustainability of sweet cherry production in Europe and provide a case for breeders to develop cultivars adapted to the upcoming climatic conditions. This is especially pertinent for species for which no low-chill cultivars have been bred so far, such as sweet cherry and apricot.

Adaptation options other than new cultivars or new growing regions are currently available to growers. In particular, dormancy breaking agents can be used to compensate for insufficient chilling (Erez 2000). Greater understanding of these agents’ ability to compensate for insufficient chill under climate change is required to understand if these provide an enduring adaptation strategy. Reliance on these products may not be a sustainable option with regulations in recent years restricting product allowable (e.g. Dormex®).

These projections were made using two GCMs. Ideally, a full ensemble of models would be used to assess potential impact (e.g. CMIP 5; Taylor et al. 2012)). Here, we used two models among the limited set available with data for this assessment (projections with daily temperature data). For the purposes of this assessment, this was sufficient to illustrate changes in risk based on CR definitions and a greater set of scenarios may be used to further explore the risk analysis. Furthermore, this analysis used gridded temperature data that may not always be representative of microclimates within orchards and as such, thus these risk projection results should be considered a guide only.

### Climate change: a shift in growing areas and cultivars

For the most pessimistic climate scenario (Fig. 4), the projections show notable risk for chill fulfilment of high chill cultivars in areas currently considered as sufficiently cold, i.e. Greece, Italy, France and British Isles. This highlights that high chill cultivars will need to be replaced with lower chill cultivars. However, these cultivar replacements need to be considered within the wider agri-climate setting. For example, cultivars with lower chill requirements tend to be characterised by earlier flowering dates, especially under warming spring temperatures and may therefore be at higher risk of early-season frost damage (Bigler and Bugmann 2018; Vitasse et al. 2018). Selection of appropriate cultivars therefore require a wider risk assessment to avoid maladaptation.

Interestingly, new areas suitable for cultivars with relatively high chill requirements appear in North-Eastern Europe by the end the century. This result stems from experiments which indicated temperatures below 0 to −2°C are not effective for chill accumulation as they might inhibit all cellular functions and this low temperature restriction has been included in most models of chil (e.g. (Richardson et al. 1974; Fishman et al. 1987; Hänninen 1990). Consequently, current climatic conditions that are too cold to satisfy chill requirements, according to current understanding of chill accumulation, will improve in suitability for chill accumulation and illustrates new production areas suitable for fruit tree cultivation. Again, other agri-climate risks (e.g. frost risk, water availability) need to be evaluated prior to establishing new growing regions.

Overall, this paper reveals that the method used to assess chilling requirements does have an important impact on assessments of the regional and cultivar suitability of sweet cherry. To more broadly understand the risk warming will pose to crops that need a high bloom load for profitability we propose the evaluation of chilling requirements using higher percentages of bud break (around 90%) over a longer period under forcing conditions. This approach should be undertaken to assess chilling requirements for the evaluation of new cultivars. These findings can help inform selections made by breeders to develop climate adapted cultivars and for growers to better understand current and future production risk in current and new growing areas.

## Acknowledgements

The authors would like to acknowledge the E-OBS dataset from the EU-FP6 project ENSEMBLES (http://ensembles-eu.metoffice.com) and the data providers in the ECA&D project (http://www.ecad.eu). JAC was supported by CEP Innovation and Aquitaine Region (AQUIPRU project 2014-1R201022971). The authors warmly thank Teresa Barreneche, Hélène Christmann, Jacques Joly, Lydie Fouilhaux, Noémie Vimont and Rémi Beauvieux for collecting the branches and collaborating on the phenotyping. The authors thank the INRA’s ‘*Prunus* Genetic Resources Center’ for preserving and managing the sweet cherry collections and the Fruit Experimental Unit of INRA-Bordeaux (UEA) for growing the trees and managing the orchards. Finally, many thanks to Alexis Berg for his help on the analysis of climatic projection data.

